# Analysis of immunological resistance to primary *Mycobacterium tuberculosis* infection in humans

**DOI:** 10.1101/551432

**Authors:** January Weiner, Teresa Domaszewska, Simon Donkor, Philip C. Hill, Jayne S. Sutherland

**Affiliations:** Max Planck Institute for Infection Biology, Berlin, Germany; Vaccines and Immunity Theme, Medical Research Council Unit The Gambia at London School of Hygiene and Tropical Medicine, Banjul, The Gambia; Otago University, Otago, New Zealand

## Abstract

**Background:** Despite recent advances in diagnosis and treatment, tuberculosis (TB) remains a major infectious disease killer in resource-poor settings. Strategies to prevent *Mycobacterium tuberculosis* (Mtb) infection are urgently required. By characterising natural protective immunity to Mtb infection we aimed to identify correlates of protection to guide vaccine development and other immune based therapies.

**Methods:** Two groups of Mtb-exposed contacts of TB patients were recruited in The Gambia and assessed for Mtb infection status using either tuberculin skin test (TST) reactivity at baseline and 3 months or QuantiFERON (QFT) reactivity at baseline and 6 months. For both groups, converters were defined as having a negative test at baseline and a positive one at follow-up, while those with a negative test at both time-points were defined as non-converters (Mtb resisters). Participants were analysed using RNA-sequencing and plasma Mtb proteome IgA and IgG arrays.

**Results:** Several genes were found to be differentially expressed at baseline between the groups prior to any signs of infection by current tests. Modular analysis revealed a distinct B cell gene signature in TST non-converters compared to converters (at q < 10^-6^, AUC > 0.7), which was only present in the most highly exposed group. Interestingly, when infection status was defined by QFT, enrichment of Type I IFN and antiviral gene signatures was observed. Plasma IgG and IgA antibody reactivity across the entire Mtb proteome showed the best differentiation in individuals with the highest exposure. An AUC of 1.0 (q<10^-3^) was observed for IgA reactivity to Rv0134 and an AUC of 0.98 for IgA reactivity to both Rv0629c and Rv2188c (all lower in TST non-converters). IgG reactivity to Rv3223c resulted in an AUC of 0.96 (q < 10^-4^) and was again lower in TST non-converters. The highest AUC for those with lower Mtb exposure were 0.84 (Rv2411c) for IgA and 0.83 (Rv2131c) for IgG.

**Conclusions:** These data provide insight into the early protective response to Mtb infection and possible avenues for novel therapeutic strategies to prevent Mtb infection.

## Introduction

Despite recent advances in diagnosis and treatment, tuberculosis (TB) remains is the leading infectious disease killer globally. Over 2 billion people are latently infected with *Mycobacterium tuberculosis* (Mtb) with 10 million new cases and 1.6 million deaths each year [1]. Strategies to prevent Mtb infection occurring in the first place are a high priority. A recent study showed that sustained QuantiFERON (QFT) conversion, which may best reflect the establishment of Mtb infection, was reduced by 45.4% with Bacillus Calmette–Guérin (BCG) re-vaccination of adolescents in a high-transmission setting [2]. An understanding of the underlying mechanism(s) for protection against infection are required in order to inform clinical development of new vaccine candidates and host-directed therapies.

The World Health Organization (WHO) strategy sets ambitious targets to ‘End TB’ through a multi-pronged attack, including a major escalation of research activities to identify novel approaches [3]. Therapeutics to prevent/clear Mtb infection is an alternative strategy that could turn the tide on the TB epidemic. Evidence for natural resistance to Mtb infection is best illustrated by the Lübeck disaster where ∼20% of infants vaccinated with Mtb-contaminated BCG did not get infected [4]. In addition, studies of sailors in long-term confinement with a TB patient showed a surprisingly low level of infection [5]. It has also been shown that a proportion of highly Mtb-exposed household contacts never acquire Mtb infection [6]. Importantly, longitudinal studies of such individuals confirm a much lower rate of progression to TB disease than those test positive for LTBI [7] suggesting that they are protected and not merely anergic to the antigens in diagnostic tests.

Several cell subsets have been proposed to control resistance to Mtb infection including macrophages, non-classical T cell subsets and B cells. Macrophage-dependent pathways that prevent bacterial uptake or rapidly clear Mtb before the development of an adaptive immune response define innate resisters while adaptive resisters are individuals in which T cell and B cell effector functions eliminate or restrict Mtb infections, either independently of IFN-γ production or through priming by non-protein antigens [8]. Further study is required to determine whether adaptive resisters have greater resistance to progression to active TB than individuals with traditionally defined LTBI [8]. Genome-wide association analysis has also identified loci that are associated with innate or adaptive resistance to Mtb infection [9].

The aim of this study was to analyse transcriptomic and antibody signatures in Mtb infection converters and non-converters to identify subsets and pathways for rational development of novel interventions to enhance protection against Mtb infection.

## Methods

### Study participants

Household contacts of TB index patients were recruited following written, informed consent by the participant or a parent/guardian if <18 years. They were screened to rule out active disease and infection status was determined using either Tuberculin skin test (TST) or QuantiFERON (QFT; Qiagen). TST was performed for participants recruited between 2002-2005 at baseline and 3 months using 2 tuberculin units (TU) of purified protein derivative (PPD) RT23 (SSI, Denmark) with a reading ≥10mm considered positive. QFT (Qiagen, Germany) was performed for participants recruited between 2016 and 2017 at baseline and 6 months according to the manufacturer instructions. Two participant groups were predefined: TST converters (those who were negative (0mm) at baseline and converted to positive (≥10mm) by 3 months) and TST non-converters (those who remained 0mm at both time-points). Similarly, analysis of QFT at baseline and 6 months defined QFT converters and non-converters. Whole blood RNA was stabilised in Paxgene RNA blood tubes and stored at −80 until analysis. Heparinised blood was centrifuged (600_gmax_, 10 mins.), the plasma collected and stored at −20°C until analysis. Differential blood counts were performed at baseline for all participants using a hematology analyser (Cell-Dyn, USA).

### RNA-sequencing

RNA was extracted (Qiagen, Germany) and shipped to the Beijing Genome Institute (Hong Kong) and underwent Globin transcript depletion (GlobinClear, Life Technologies, UK). cDNA libraries were prepared using Illumina mRNA-Seq Sample Prep Kit and RNA-Seq was performed by Expression Analysis Inc., at 20 million 50bp paired-end reads, on Illumina HiSeq-2000 sequencers. Read pairs were aligned to the hg19 human genome using gsnap24 which generated splice junction counts for each sample.

### 4000 Mtb proteome arrays

IgG and IgA serum antibody reactivity to 4000 Mtb antigens was analysed using a full proteome microarray by Antigen Discovery Inc. (USA) as previously described [10, 11]. Briefly, proteome chips were probed with serum from TST converters and non-converters at baseline and 3 months. Slides were first blocked for 30 min in protein array-blocking buffer before incubation with the primary antibody for 2h. Antibodies were detected with Cy3-conjugated secondary antibodies (Jackson ImmunoResearch) and scanned in a ScanArray 4000 laser confocal scanner (GSI Lumonics, Billerica, MA). Fluoresence intensities were quantified by using QUANTARRAY software (GSI Lumonics).

## Statistics

### Participant demographics

Data were analysed a using Mann-Whitney U-test or Kruskal-Wallis test with Graph Pad Prism software v7.0 (Software MacKiev, USA).

### RNA-sequencing

RNA Seq count data were processed using the edgeR package. For each batch, only genes with at least 5 counts per million in at least 3 samples were kept. For batch 1, a subset of samples was generated comprising only samples from HIV- individuals older than 17 years. Differential gene expression was analysed using glmLRT function from the edgeR package, which is a log-likelihood ratio test. Discordance analysis was performed using the disco package. Gene set enrichment analysis was performed using the tmod package. In addition, due to subset differences within each batch the ComBat function from the sva R package was used to remove the effect of sub-batches from logarithmized counts per million transformed data.

### Antibody arrays

For each antigen, the average intensity of all samples was determined. A Mann-Whitney U-test was performed for analysis between groups and adjusted for false discovery rates using the Benjamini-Hochberg test. The Area under the ROC curve (AUC) was used to determine differential reactivity for each antigen for both IgG and IgA.

## Results

### Participant demographics

For metabolomics and antibody arrays, paired plasma samples from 40 converters and 38 non-converters at baseline and 3 months were sent to Metabolon and Antigen Discovery for processing respectively. No difference in age or sex was seen between the groups but there was a significantly higher proportion of converters living in the same room as the index case (ie closest proximity; p=0.034) compared to non-converters (**Table 1**). For RNA-sequencing, analysis was performed on two batches based on TST (batch 1) or QFT (batch 2). For batch 1, 100 baseline RNA samples were sent to BGI for sequencing. Quality control prior to library preparation showed that only 46/100 samples were of optimal quality for sequencing (RIN≥7.0) presumably due to the long storage duration (up to 13 years). Of these 46 samples, 35 were TST non-converters and 11 were TST converters (**Table 1**). There was a similar proportion of males in both groups, but a significant difference in age (median[IQR] non-converters = 10[5-17] compared to 20[16-32] for the converters (p<0.001). There was no significant difference in proximity between the groups. For batch 2, samples from 16 QFT non-converters and 11 QFT converters were sequenced (**Table 1**). No significant difference in age, sex or proximity was seen between the groups. Progression to active disease was assessed in all participants for a 24 month period with only 1 QFT converter progressing after 12 months and none of the non-converters progressing. No progressors were seen in the TST converter/non-converter group.

**Table 1:** Participant demographics:

|  | **Non-converters** | **Converters** | **p-value** |
| --- | --- | --- | --- |
| **Plasma** | **n=40** | **n=38** |  |
| Age (median[IQR]) | 22[18-34] | 26[20-37] | ns |
| Males (n(%)) | 14 (35) | 14 (37) | ns |
| **Proximity (n(%))** |  |  |  |
| Different house | 15 (38) | 10 (26) | ns |
| Different room | 21 (53) | 17 (45) | ns |
| Same room | 4 (10) | 11 (29) | 0.034 |
| **RNA TST** | **n=35** | **n=11** |  |
| Age (median[IQR]) | 10[5-17] | 20[16-32] | <0.001 |
| Males (n(%)) | 18 (51) | 5 (45) | ns |
| **Proximity (n(%))** |  |  |  |
| Different house | 19 (54) | 6 (55) | ns |
| Different room | 14 (40) | 5 (45) | ns |
| Same room | 3 (9) | 2 (18) | ns |
| **RNA QFT** | **n=16** | **n=11** |  |
| Age (median[IQR]) | 29[19-47] | 18[17-34] | ns |
| Males (n(%)) | 8 (50) | 4 (36) | ns |
| **Proximity (n(%))** |  |  |  |
| Different house | 0 | 0 | ns |
| Different room | 0 | 0 | ns |
| Same room | 16 (100) | 11 (100) | ns |

### RNA-sequencing

Several differentially expressed genes were observed between converters and non-converters at baseline. For the TST group, the most differentially expressed genes included TMEM56 (p=1.25E-08), RP11-364L4.1 (p=4.52E-08), LCN2 (p=1.81E-07), CLIC2 (p=3.50E-07) and WASHC3 (p=3.54E-07) (**Table 2**). For the QFT group, the most differentially expressed genes included RPS9 (p=3.14E-10), MDM4 (p=6.09E-07), HSFX3 (7.6E-07) and IGF1R (p=5.25E-06) all of which were upregulated in the converters (**Table 3**). A full description of differentially expressed genes is shown in **supplementary table 1**. Due to the significant difference in age between the TST converters and non-converters, we also performed filtered analysis of adults only (**Table 4**). Interestingly in this group the top genes included several coding for immune-regulatory functions including CXCL10 (p=3.10E-05), HLA-DQB1 (p=8.53E-04) and CD22 (p=1.58E-03). However, significance was lost after adjusting for FDR, most likely due to low numbers in both groups. CXCL10 (IP-10) showed a significant down-regulation in the non-converters while CD22 (inhibitory receptor for B cell receptor signalling; member of Ig superfamily) showed an up-regulation. Interestingly, HLA-DQB1 was highly upregulated in the TST non-converters (logFC = −5.6) suggesting this gene may be involved in protection against Mtb infection. Next we performed gene set enrichment analysis using the Tmod package in R. In the total TST-defined group, there was a strong enrichment in B-cell related genes in the TST non-converters **(Fig. 1A)**, which was still evident in the filtered subgroup (**Fig. 1B**). TST converters had significant enrichment for TLR8-BAFF related genes and monocytes (**Fig. 1A**) although this was lost when filtering was performed. The responses in the QFT non-converters were dominated by an interferon-Type I/anti-viral gene signature (**Fig. 1C**). The B cell genes incorporated into M47.0, M47.1 and M69 include BLK, CD19, CD22, CD24, CD72, CD79, CD200, CXCR5, CR2 and FCR1/2. The genes that contribute to the Type I IFN response module (LI.M127) include TAP1, IFIH1, IRF7, PARP9, STAT1, PLSCR1, IFITM1, HERC5, DDX60, USP18, RSAD2 and IFIT1.

**Table 2.**
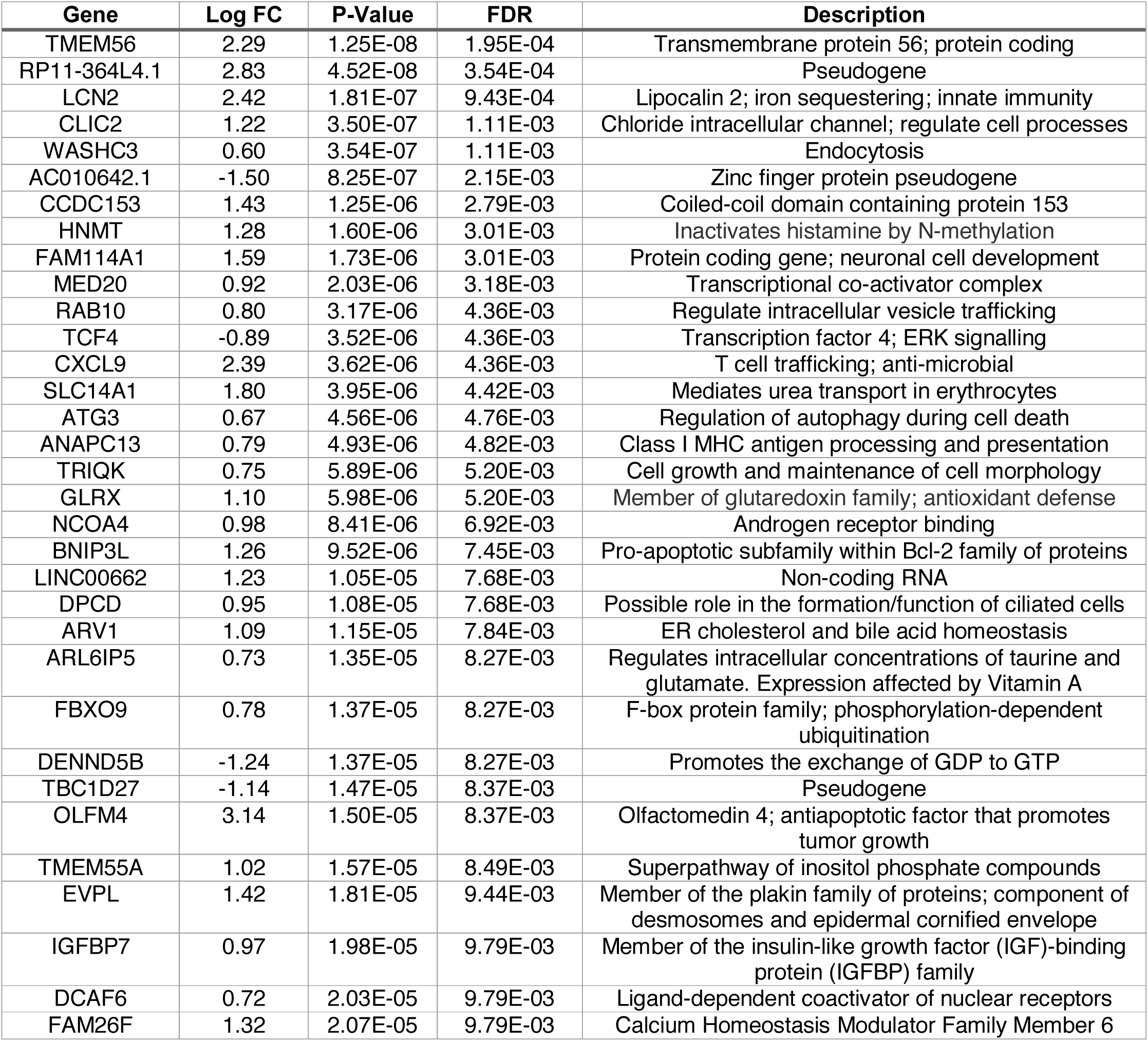
Differentially expressed genes in the TST group with FDR ≤0.01.

**Table 3.** Differentially expressed genes in the QFT group with FDR ≤0.05.

| **Gene** | **Log FC** | **P-Value** | **FDR** | **description** |
| --- | --- | --- | --- | --- |
| RPS9 | 5.39 | 3.14E-10 | 4.87E-06 | ribosomal protein S9 |
| MDM4 | 0.73 | 6.09E-07 | 3.97E-03 | MDM4, p53 regulator |
| HSFX3 | 6.04 | 7.66E-07 | 3.97E-03 | heat shock transcription factor family |
| RP60S | -0.92 | 4.10E-06 | 1.59E-02 | 60S ribosomal protein pseudogene |
| IGF1R | 0.72 | 5.25E-06 | 1.63E-02 | insulin like growth factor 1 receptor |
| STRN3 | 0.80 | 9.23E-06 | 1.80E-02 | striatin 3; calmodulin binding protein |
| RPL39P3 | -0.85 | 9.27E-06 | 1.80E-02 | ribosomal protein L39 pseudogene 3 |
| PCNA | -0.63 | 9.60E-06 | 1.80E-02 | proliferating cell nuclear antigen |
| PCK2 | -0.62 | 1.18E-05 | 1.80E-02 | mitochondrial |
| POLR2L | -0.63 | 1.26E-05 | 1.80E-02 | RNA polymerase II subunit L |
| TFP1 | 5.47 | 1.31E-05 | 1.80E-02 | transferrin pseudogene 1 |
| SYNE3 | 0.59 | 1.39E-05 | 1.80E-02 | Actin binding |
| LAMTOR2 | -0.68 | 1.71E-05 | 1.82E-02 | MAPK and MTOR activator 2 |
| RNF181 | -0.60 | 1.73E-05 | 1.82E-02 | ring finger protein 181 |
| NKTR | 0.73 | 1.82E-05 | 1.82E-02 | natural killer cell triggering receptor |
| PMVK | -0.60 | 1.87E-05 | 1.82E-02 | phosphomevalonate kinase |
| C19orf70 | -0.60 | 2.07E-05 | 1.89E-02 | chromosome 19 open reading frame 70 |
| DPM3 | -0.67 | 2.18E-05 | 1.89E-02 | dolichyl-phosphate mannosyltransferase |
| HLA-DMA | -6.98 | 2.51E-05 | 2.06E-02 | MHC class II, DM alpha |
| HLA-B | -6.14 | 2.90E-05 | 2.13E-02 | MHC class I, B |
| HNRNPH1 | 0.65 | 2.96E-05 | 2.13E-02 | heterogeneous nuclear ribonucleoprotein H1 |
| RARRES3 | -0.67 | 3.02E-05 | 2.13E-02 | retinoic acid receptor responder 3 |
| RBM25 | 0.58 | 4.35E-05 | 2.74E-02 | RNA binding motif protein 25 |
| LILRB3 | -3.44 | 4.46E-05 | 2.74E-02 | leukocyte immunoglobulin like receptor B3 |
| TMSB10 | -0.59 | 4.54E-05 | 2.74E-02 | thymosin beta 10 |
| STX16 | 0.48 | 4.61E-05 | 2.74E-02 | syntaxin 16 |
| FBXO6 | -0.80 | 4.77E-05 | 2.74E-02 | F-box protein 6 |
| MALAT1 | 0.69 | 5.26E-05 | 2.90E-02 | metastasis associated lung adenocarcinoma |
|  | 0.87 | 5.47E-05 | 2.90E-02 | novel transcript, sense intronic to ST20 |
| RPL29P11 | 2.33 | 5.60E-05 | 2.90E-02 | ribosomal protein L29 pseudogene 11 |
| LMBR1L | 0.47 | 6.41E-05 | 3.20E-02 | limb development membrane protein 1 like |
| TMEM170B | 0.80 | 6.59E-05 | 3.20E-02 | transmembrane protein 170B |
| LTB | 5.44 | 7.12E-05 | 3.35E-02 | lymphotoxin beta |
| PLEKHF1 | -0.69 | 7.33E-05 | 3.35E-02 | triggers caspase-independent apoptosis |
| SERPING1 | -1.76 | 7.85E-05 | 3.49E-02 | complement component 1 inhibitor |
| LINC01089 | 0.67 | 8.31E-05 | 3.59E-02 | long intergenic non-protein coding RNA 1089 |
|  | 0.96 | 9.11E-05 | 3.83E-02 | uncharacterized LOC100288123 |
| SF3B1 | 0.49 | 9.67E-05 | 3.87E-02 | splicing factor 3b subunit 1 |
| NDUFA4 | -0.50 | 9.71E-05 | 3.87E-02 | NDUFA4, mitochondrial complex associated |
| NKG7 | -0.85 | 1.05E-04 | 4.01E-02 | natural killer cell granule protein 7 |
| ULK1 | 0.53 | 1.06E-04 | 4.01E-02 | unc-51 like autophagy activating kinase 1 |
| SRSF11 | 0.57 | 1.15E-04 | 4.14E-02 | serine and arginine rich splicing factor 11 |
| EPSTI1 | -1.18 | 1.17E-04 | 4.14E-02 | epithelial stromal interaction 1 |
| EIF4A1 | 0.98 | 1.17E-04 | 4.14E-02 | eukaryotic translation initiation factor 4A1 |
| ATP11A | 0.47 | 1.23E-04 | 4.24E-02 | ATPase phospholipid transporting 11A |
| PSMG3 | -0.57 | 1.52E-04 | 4.91E-02 | proteasome assembly chaperone 3 |
| OAS1 | -1.46 | 1.53E-04 | 4.91E-02 | 2'-5'-oligoadenylate synthetase 1 |
| SERINC5 | 0.69 | 1.54E-04 | 4.91E-02 | serine incorporator 5 |
| BANF1 | -0.57 | 1.58E-04 | 4.91E-02 | barrier to autointegration factor 1 |
| FOXK1 | 0.55 | 1.59E-04 | 4.91E-02 | forkhead box K1 |
| AQR | 0.67 | 1.61E-04 | 4.91E-02 | aquarius intron-binding spliceosomal factor |
| PAN3 | 0.45 | 1.66E-04 | 4.93E-02 | poly(A) specific ribonuclease subunit PAN3 |
| ECI1 | -0.55 | 1.68E-04 | 4.93E-02 | enoyl-CoA delta isomerase 1 |
| ZMYND15 | 1.10 | 1.73E-04 | 4.98E-02 | zinc finger MYND-type containing 15 |

**Table 4.** Differentially expressed genes in the TST group ≥15 years with FDR ≤0.05.

| Gene | Log FC | p-value | FDR | Description |
| --- | --- | --- | --- | --- |
| CXCL10 | 3.97 | 3.10E-05 | ns |  |
| MTRNR2L1 | 4.19 | 3.23E-05 | ns |  |
| LILRB2 | 3.61 | 4.67E-04 | ns |  |
| HNMT | 1.40 | 4.89E-04 | ns |  |
| EEF1A1P5 | 1.50 | 6.91E-04 | ns |  |
| ADAMTS1 | 1.51 | 7.54E-04 | ns |  |
| HLA-DQB1 | -5.65 | 8.53E-04 | ns |  |
| SRGAP3 | -1.10 | 9.82E-04 | ns |  |
| HOPX | 0.85 | 1.12E-03 | ns |  |
| CD22 | -0.84 | 1.58E-03 | ns |  |
| CLIC2 | 1.16 | 1.65E-03 | ns |  |
| RP11-394B2.5 | -0.85 | 1.70E-03 | ns |  |
| FAM26F | 1.60 | 1.71E-03 | ns |  |
| CCDC114 | -1.22 | 1.74E-03 | ns |  |
| GSKIP | 0.67 | 1.94E-03 | ns |  |
| MRPS35 | 0.60 | 2.08E-03 | ns |  |
| RP11-374P20.4 | -1.44 | 2.19E-03 | ns |  |
| LCN2 | 2.07 | 2.43E-03 | ns |  |
| AC020951.1 | -1.17 | 2.68E-03 | ns |  |
| KIF19 | -1.02 | 2.74E-03 | ns |  |

**Figure 1:**
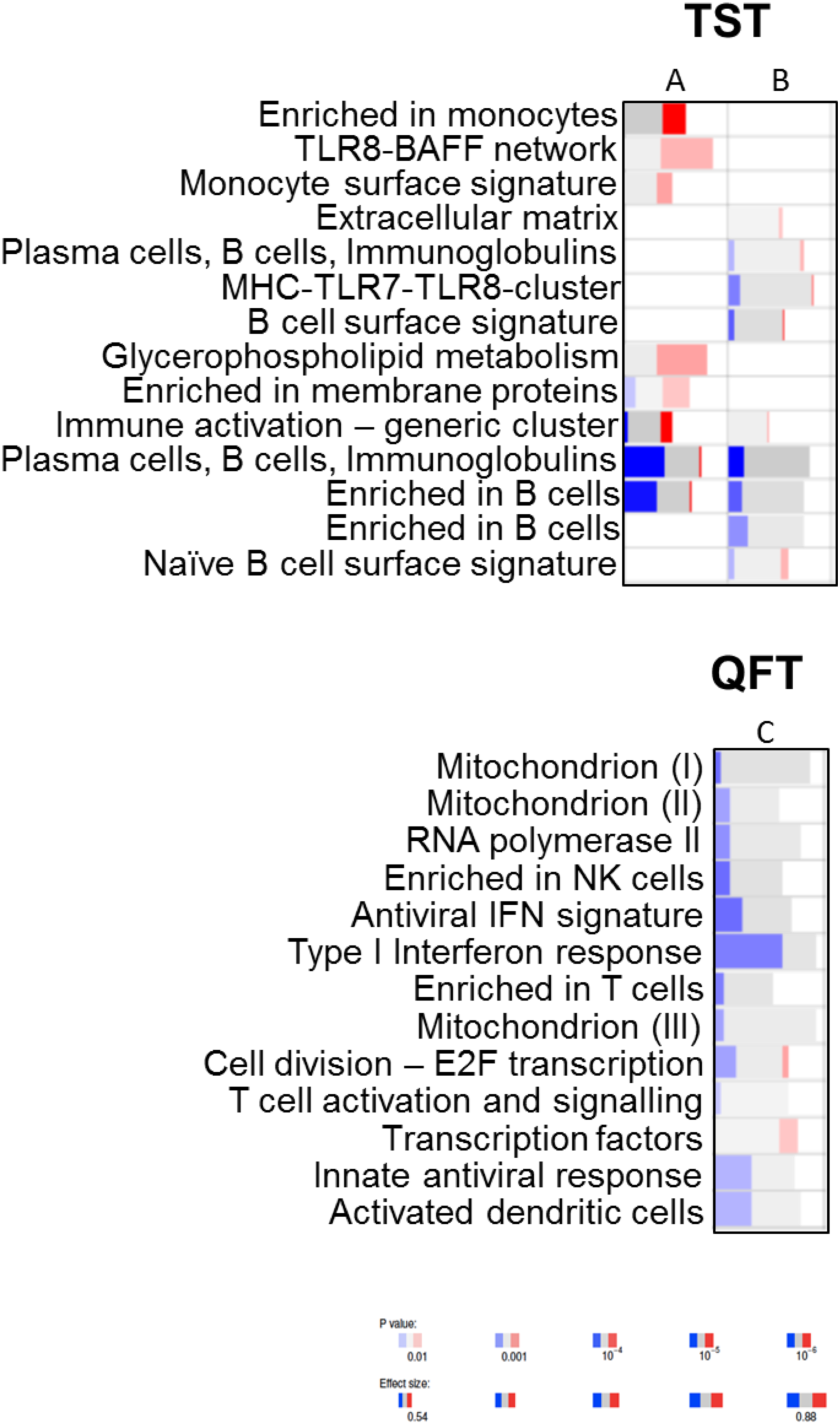
RNA-seq analysis modular enrichment of genes in TST non-converters compared to converters (A, B) or QFT non-converters compared to converters (C). Blue indicates higher expression in non-converters. Effect size (box length) indicates area under the curve (AUC).

It is important to note there were no significant differences in white blood cell counts between converters and non-converters (data not shown). However, due to the differences in proximity (ie exposure) of each participants, we next assessed how this affected gene expression profiles (**Fig. 2**). Interestingly, in those with the lowest exposure, gene signatures were generally within the innate immune cells with enrichment of neutrophils, monocytes and TLR pathways. The B cell signature was only evident in those with the highest exposure to the index TB case (**Fig. 2**). This suggests that B cells are involved in the early protective immune response to Mtb infection.

**Figure 2:**
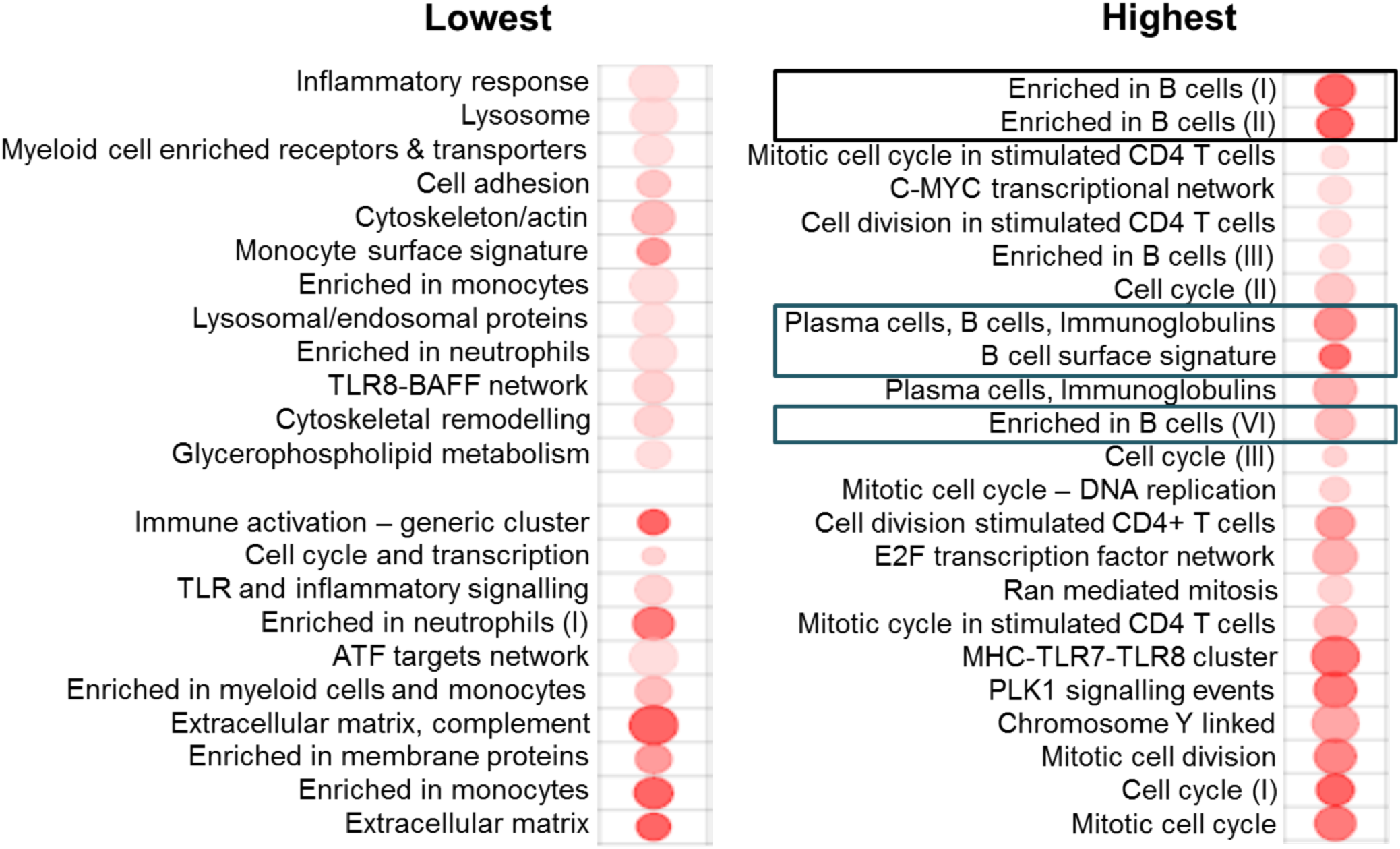
RNA-seq analysis showing module enrichment genes in those with low (left) or high (right) exposure to a TB-index case. B cell enrichment was only seen in those with the highest exposure.

### Mtb antibody arrays

Analysis of both IgG and IgA reactivity to the Mtb proteome showed similar findings with extensive differences in reactivity evident (**supplementary Table 2**). There was a significant exposure gradient effect with the AUC increasing as proximity to the index case increased (**Fig. 3**). For example, with IgG reactivity to Rv2131c, Rv0363c and Rv3223c the AUC was 0.73, 0.83 and 0.96 respectively and with IgA reactivity to Rv0134 at the closest proximity, the AUC was 1.00. This indicates that antibody reactivity can be used to discriminate between TST converters and non-converters. Surprisingly, however, the majority of reactivity was higher in the converters (**Fig. 3 (red)**) suggesting that the enrichment of B cell genes we saw in the non-converters was not primarily due to enhanced antibody production. However, when the most reactive antigens were analysed, non-converters showed higher responses to Rv0831c (fold increase (fi) 1.93), Rv3038c (fi 2.32), Rv2946c (fi 2.09), Rv3604c (fi 2.53), Rv0726c (fi 1.93) and Rv2396 (fi 2.00) (**Fig. 4**).

**Figure 3:**
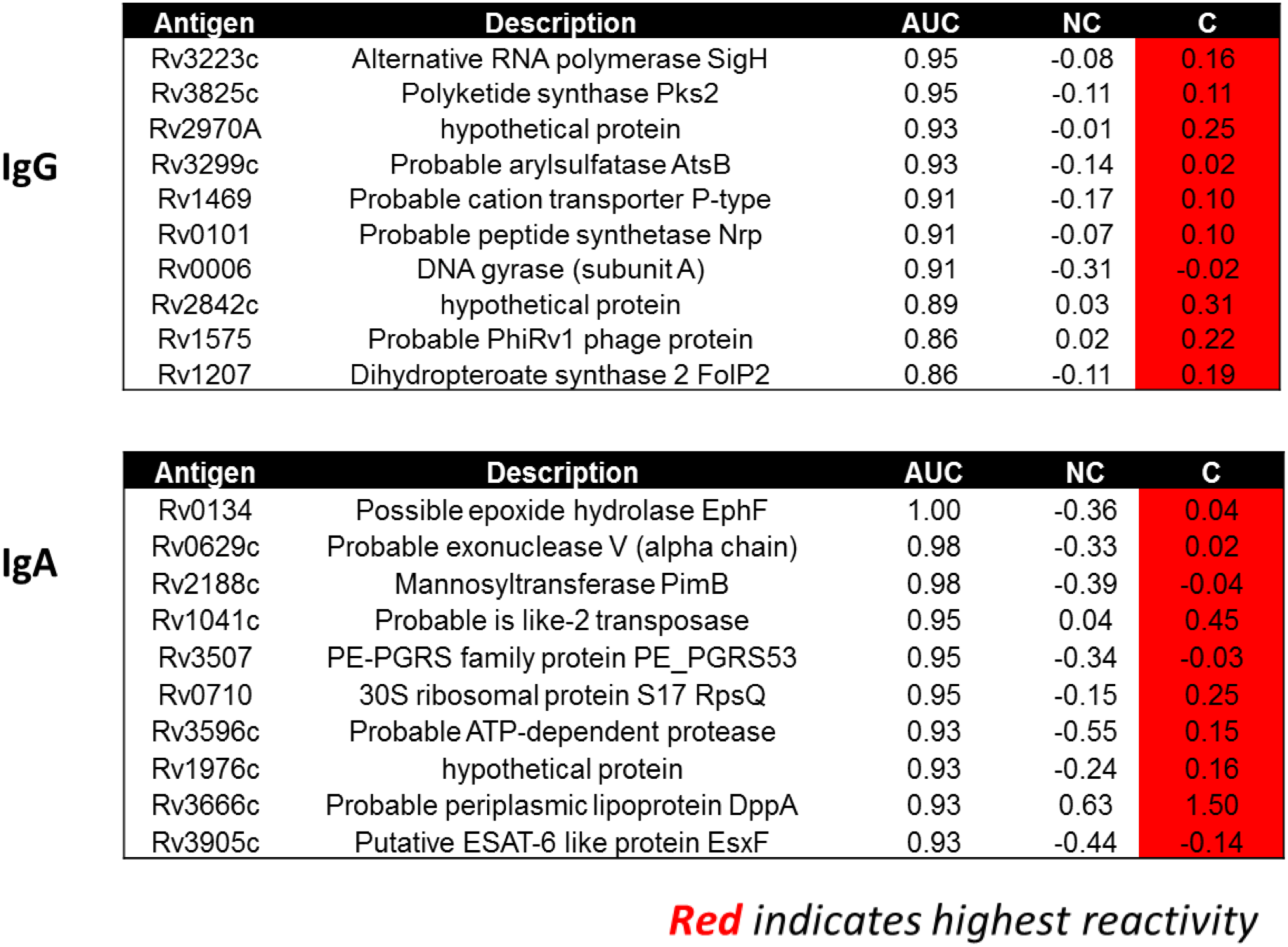
IgG (top) and IgA (bottom) reactivity to Mtb antigens in TST converters (C) and non-converters (NC).

**Figure 4:**
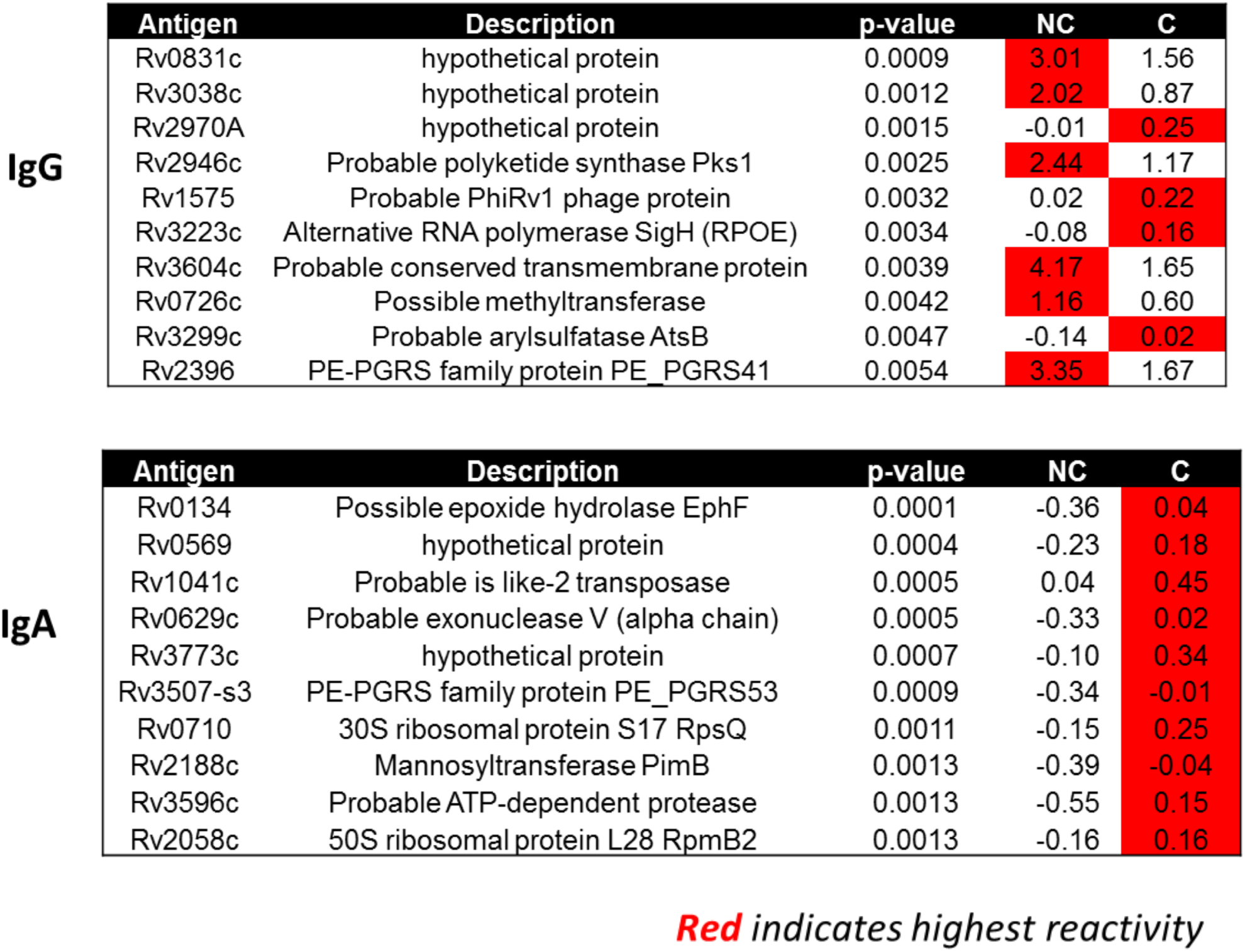
Antigens showing the greatest IgG (top) and IgA (bottom) reactivity in TST converters (C) and non-converters (NC) at baseline.

We next analysed reactivity to ESAT-6 specific antigens since this is a dominant secretory antigen from Mtb. Only individuals in the closest proximity to the index case showed any differential antibody reactivity to ESAT-6 antigens with the most significant being IgG reactivity to Rv3875 (p=0.0008). IgA reactivity to Rv3872 (p=0.025) and Rv3871 (p=0.030) were both increased in converters compared to non-converters at baseline (**supplementary Table 2**).

## Discussion

We have characterised host transcriptomic and antibody responses to Mtb in TST non-converters and converters prior to any signs of infection as determined by current methods. We have previously shown higher levels of soluble IL-17 in non-converters at baseline [12] suggesting they have encountered Mtb and mounted an immune response and are not simply anergic to antigens in the current tests nor is Mtb blocked by physiological barriers. In this study we saw a distinct B cell gene signature reflected by differential antibody responses in TST converters and non-converters to multiple Mtb antigens. Interestingly, defining infection based on QFT rather than TST showed a distinct Type I interferon/anti-viral gene signature.

Analysis of RNA from converters and non-converters at baseline highlighted several differentially expressed genes. These included CXCL9, which is a T-cell chemoattractant induced by IFN-γ [13]; and BNIP3L (BCL2 Interacting Protein 3 Like), a proapoptotic protein, which may play a role in tumor suppression [14] were both upregulated in converters. Consistent with our findings from metabolomics analysis, we also saw upregulation of SLC14A1, which is involved in urea transportation [15] and LCN2, which is involved in iron sequestration and innate immunity [16]. Both of these genes are important in the anti-microbial response and both were upregulated in converters prior to conversion. Interestingly, when modular analysis was performed, we saw a significant enrichment of B cell genes in non-converters compared to converters, whilst converters had enrichment for monocytes and TLR8-BAFF – important in the early airway epithelial response to Mtb [17]. The B cell genes of interest included CD72, which is thought to mediate B cell-T cell interaction whilst CD19, CD22, CD24 and CD200 are all phenotypic/maturation markers. Interestingly high affinity FCR1/2 genes are enriched in active TB compared to latent TB [18] and was recently shown to be a prognostic marker of progression to active TB in Gambian household contacts [19].

It was interesting to see the distinct gene signatures present when infection was defined by QFT rather than TST. Whilst different mechanisms will be inevitable since TST is based on a delayed type hypersensitivity reaction of 48-72 hours while QFT is based on overnight *in vitro* stimulation, the fact these signatures are upregulated in the uninfected groups, suggests they are likely to play a role in resistance. Increased production of Type I interferons (IFNα/β) as part of the anti-viral response inhibits the downstream effects of Type II interferon (IFN-γ) responses known to be critical for Mtb control [20]. Thus it was surprising to see an increase in these genes in the QFT non-converters. However, it has previously been shown that in the absence of a response to IFN-γ, type I IFNs play a non-redundant protective role against tuberculosis in mice [21] and suggest type I IFN can limit the number of target cells that Mtb can infect in the lungs while IFN-γ enhances their ability to restrict bacterial growth [21]. It has also been shown in experimental models that Type I IFN may play a protective role in the context of BCG-induced immunity and could be targeted to improve preventive vaccination against tuberculosis [22].

The enrichment of B cell genes was unexpected so we decided to analyse IgG and IgA plasma antibody reactivity to all 4000 Mtb proteins from these same individuals. TST converters had significantly higher levels of both IgG and IgA antibodies to multiple Mtb antigens at baseline, which allowed excellent discrimination between converters and non-converters. This was particularly evident when proximity was accounted for: exposed contacts in closest proximity to the index case who later converted their TST had higher antibody levels at baseline. This suggests that antibodies are playing a role in the response to Mtb infection but are not necessarily protecting the host from becoming infected (although a few antigens elicited significantly higher IgG responses in the non-converters). Since we saw such a distinct B cell signature in the non-converters, this suggests that other antibody-independent functions of B cells are driving the protection from infection and this is currently being assessed in our laboratory.

In conclusion, despite being clinically healthy and showing no signs of latent infection by current tests, we were able to show differential immune responses in subjects who would later convert to a positive TST compared to those who remained TST negative. Our results support the recent proposed criteria to define resistance by Simmonds et al [8], including the use of both TST and QFT with multiple negative results in the first 12 months following exposure. The interesting B cell and Type I IFN signatures provide an avenue for analysis of mechanisms underlying the protection and could be targeted to enhance resistance to Mtb.

## Acknowledgements

We would like to thank the National TB control program, participants and their families. We also thank the MRC Gambia TB Clinic staff, TB immunology and TB microbiology laboratory staff. The clinical cohort was funded through the MRC Gambia core funding. All experimental analysis was funded through a TBVAC2020 sub-grant awarded to A/Prof Sutherland (Grant number SEP-210138189).

